# Spatially Anchored Regulatory State Inference in Melanoma

**DOI:** 10.64898/2026.04.05.716552

**Authors:** Jagan Mohan Reddy Dwarampudi, Veena Kochat, Suresh Satpati, Md Ishtyaq Mahmud, Humaira Anzum, Khalida Wani, Alexander Lazar, Ajay Kumar Saw, Jared Malke, Hien V. Nguyen, Kunal Rai, Tania Banerjee

## Abstract

Spatial transcriptomics (ST) captures gene expression within tissue architecture but lacks direct regulatory information, while single-cell multiome assays profile transcriptional and chromatin states without spatial context. We present a framework for spatially anchored regulatory inference that integrates Visium ST with single-cell multiome data to infer spatially resolved regulatory programs. Building upon GraphST, we introduce spatially regularized cell-to-spot mapping and propagate chromatin accessibility and transcription factor motif activity into tissue space. Regulatory analysis is performed at the spatial domain level via joint differential expression and accessibility testing, along with quantitative concordance assessment. Applied to melanoma tissue sections, the framework reveals spatially localized regulatory programs and shows that assignment strategy substantially affects downstream regulatory stability. This modular approach enables interpretable gene-, peak-, and transcription factor–level outputs for multimodal spatial analysis.

## 1 Introduction

Spatial transcriptomics (ST) profiles gene expression within intact tissue, linking transcriptional states to anatomical context. Sequencing-based platforms such as 10x Visium provide genome-wide coverage but at multi-cell resolution, whereas higher-resolution technologies improve spatial granularity at the cost of signal sparsity or limited gene panels. In contrast, single-cell RNA and multimodal assays achieve genome-wide single-cell profiling without spatial information.

These complementary modalities motivate methods for (i) spatial domain identification, (ii) multi-slice integration with batch correction, and (iii) cross-modal mapping of dissociated single-cell data back to tissue coordinates. While substantial progress has been made in spatial clustering and slice integration, mechanistic regulatory interpretation of spatial organization remains limited. To address this gap, we develop a computational framework for regulatory interpretation of spatially organized tissue compartments.

### Spatially informed clustering

Early approaches relied solely on transcriptomic similarity (e.g., Seurat, Louvain), often producing fragmented domains. Subsequent methods incorporated spatial priors through graphical models (Giotto [1], BayesSpace [8]), graph convolutional networks (SpaGCN [4]), or graph attention autoencoders (STAGATE [3]). GraphST [5] introduced graph self-supervised contrastive learning to enforce similarity among neighboring spots, improving boundary delineation and embedding stability. Although these approaches improve spatial domain identification, they focus primarily on transcriptomic structure and are not designed for multimodal regulatory inference.

### Cross-modal spatial integration

Recent frameworks map single-cell modalities onto ST data. SIMO [6] performs probabilistic cell-to-spot mapping sequentially. SSpMosaic [7] aligns datasets using shared gene programs. MaxFuse [2] iteratively refines weak cross-modal correspondences. Garfield constructs heterogeneous molecular–spatial graphs with variational contrastive learning [9]. While these methods advance multimodal alignment and deconvolution, they primarily emphasize mapping accuracy and transcriptome-level integration. Explicit spatial regularization of assignment matrices and systematic evaluation of downstream regulatory stability remain underexplored. Furthermore, most approaches focus on transcriptome-to-transcriptome alignment and do not explicitly integrate single-cell chromatin accessibility (scATAC-seq), limiting their capacity for regulatory interpretation of spatial organization.

Building upon the GraphST framework, we make the following contributions:

1. Augment the GraphST loss with a within-cluster smoothness regularizer on the cell-to-spot mapping matrix to enforce spatially coherent assignments;
2. Propagate single-cell chromatin accessibility and transcription factor motif activity into tissue space for spot-level regulatory profiling;
3. Perform domain-level joint differential analysis of gene expression and projected accessibility with quantitative regulatory concordance; and
4. Systematically evaluate soft, hard, and sparsified mapping strategies in terms of spatial coherence, signal retention, and downstream regulatory robustness.

Experiments demonstrate that spatial smoothness regularizer on the cell-to-spot mapping matrix improves assignment coherence and strengthens expression–accessibility concordance relative to the original GraphST formulation.

## 2 Proposed Method

We integrate paired 10x Genomics Visium spatial transcriptomics and single-cell multiome (scRNA + scATAC) data from the same patient to infer spatially anchored regulatory programs.

Let *X*_*sp*_ ∈ ℝ^*S×G*^ denote Visium gene expression (spots × genes), *X*_*sc*_ ∈ ℝ^*C×G*^ denote multiome gene expression (cells × genes), and *A*_*sc*_ ∈ ℝ^*C×P*^ denote single-cell chromatin accessibility (cells × peaks). Each Visium spot has known spatial coordinates.

Our goal is to learn a mapping matrix *W* ∈ ℝ^*S×C*^ that transfers regulatory information from single cells to spatial locations, enabling the projection of accessibility and motif activity profiles into tissue space while preserving spatial coherence.

Gene expression matrices from both modalities are preprocessed using Seurat v3 highly variable gene selection (*n*_*top*_ = 3000), library-size normalization (target sum 10^4^), log-transformation, and scaling (zero_center=False, max_value=10). The shared feature space is defined as the intersection of highly variable genes across modalities, and only in-tissue Visium spots are retained for downstream analysis.

### 2.1 Graph-based embedding and spatially-regularized mapping

#### Notation

Let *S* denote the number of Visium spots and *C* the number of single-cell multiome cells. We restrict both modalities to a shared gene set of size *G*, yielding preprocessed expression *X*_sp_ ∈ ℝ^*S×G*^ (spot expression) and *X*_sc_ ∈ ℝ^*C×G*^ (single-cell expression) as defined above. *H*^sp^ ∈ ℝ^*S×G*^ and *H*^sc^ ∈ ℝ^*C×G*^ denote the learned embeddings of *X*_sp_ and *X*_sc_, produced by the GraphST GNN (for spots) and auto-encoder (for cells), respectively. The model learns a nonnegative mapping matrix *W* ∈ ℝ^*S×C*^ with column-normalization 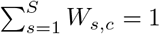 for each cell *c*, so that *W*_*s,c*_ is the fraction of cell *c* assigned to spot *s*. Spots are clustered via Leiden on the learned embedding, yielding cluster labels *ℓ*_*s*_ ∈ {1, …, *K*}. Let 𝒩_*i*_ denote the spatial neighbors of spot *i* in the *k*-NN graph. Single-cell chromatin accessibility is represented as *A*^sc^ ∈ ℝ^*C×P*^ (cells × peaks).

#### Mapping matrix and cross-modal projection

The predicted spot representation is 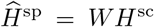. Given *W*, spot-level chromatin accessibility is projected as 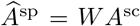. For spot-centric normalisation we define *P*_*s,c*_ = *W*_*s,c*_*/* ∑ _*c ′*_ *W*_*s,c′*_. Motif activity matrices are projected analogously.

#### GraphST mapping loss (unchanged)

The mapping matrix is learned by minimizing a loss that combines reconstruction and contrastive terms. The reconstruction loss aligns predicted and reconstructed spot representations:

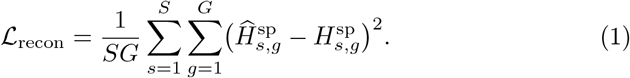

The contrastive (NCE) term enforces spatial consistency: for each spot *i*, spatial neighbors *j* ∈ 𝒩_*i*_ are positive pairs; all other spots are negatives. Let 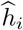 and *h*_*j*_ denote the *i*-th and *j*-th rows of 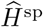 and *H*^sp^, and sim(·,·) cosine similarity.

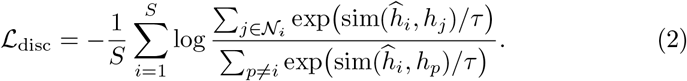

#### Spatial smoothness (our contribution)

We extend GraphST with a within-cluster smoothness regularizer on *W*. For each Leiden cluster 𝒞_*k*_ = {*s* : *ℓ*_*s*_ = *k*} with *n*_*k*_ = |𝒞_*k*_| ≥ 2, we penalize the variance of each column of *W* over spots in that cluster:

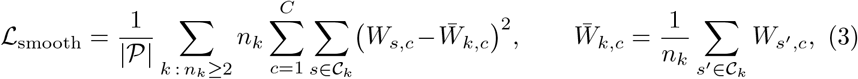

where 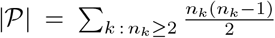 is the total number of unordered spot pairs within clusters.

#### Total objective

The model is trained by minimizing

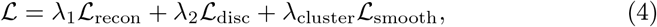

with *λ*_1_ = 10, *λ*_2_ = 1, and *λ*_cluster_ ≥ 0 controlling the strength of spatial regularization.

#### Relation to GraphST

Our framework builds upon the GraphST architecture, retaining its dual-branch embedding design, contrastive alignment strategy, and soft assignment mapping. We extend GraphST by (i) introducing a within-cluster smoothness regularizer on the mapping matrix to enforce spatially coherent assignments, (ii) propagating chromatin accessibility and motif activity through the learned mapping to obtain spot-level regulatory profiles, and (iii) systematically evaluating assignment strategies and stability across random seeds. Thus, while the embedding backbone follows GraphST, our contribution lies in spatially regularized mapping and domain-level regulatory inference.

### 2.2 Domain-Level Regulatory Inference

Spatial domains are defined by clustering of Visium spots in the learned embedding space. All downstream statistical analyses treat spots as the unit of observation.

For each domain, differential gene expression and differential chromatin accessibility are computed using Wilcoxon rank-sum tests (one-versus-rest) with Benjamini–Hochberg correction (FDR ≤ 0.05). Accessibility statistics are performed on spot-level peak matrices obtained by linear projection of single-cell ATAC profiles via the learned mapping.

To quantify regulatory concordance, we compute Pearson and Spearman correlations between domain-level log fold-changes in gene expression and corresponding gene-proximal accessibility (defined by promoter proximity to the transcription start site).

Transcription factor (TF) motif activities, computed at the single-cell level, are projected to spots through the mapping matrix and tested for domain enrichment using the same statistical framework (FDR ≤ 0.05).

Domain-specific cell-type composition is estimated by aggregating one-hot encoded cell labels through the mapping matrix, yielding spot-level cell-type proportions that are subsequently summarized per domain.

## 3 Experiments

### 3.1 Data and Experimental Setup

We analyzed paired 10x Genomics Visium spatial transcriptomics and 10x Chromium Single Cell Multiome (scRNA+scATAC) melanoma samples from four patients, comprising seven tissue sections spanning pre- and post-treatment conditions. After quality control, Visium sections contained 4,992 in-tissue spots, and multiome samples contained 8,005–16,936 cells. Shared features were defined as the intersection of highly variable genes across modalities (367–870 genes per sample).

Visium spots were connected using a *k*-nearest neighbor spatial graph (*k* = 6). Models were trained for 600 epochs using Adam with a fixed random seed unless otherwise specified.

Evaluation focuses on two aspects: (1) mapping-level behavior (sparsity, spatial coherence, stability), and (2) domain-level regulatory inference (DE–DA concordance and TF enrichment).

### 3.2 Assignment Strategy Comparison

We compared three post-hoc sparsification strategies :

– Soft: dense mapping matrix as learned.
– Hard: per-cell argmax assignment.
– Top-*k*: retain the top *k*% highest-weighted spots per cell and renormalize.

Table 1 summarizes mapping sparsity on the index section (1-Pre). Hard assignment reduces the number of non-zero mapping entries by more than four orders of magnitude but collapses the distributed contribution of cells per spot. In contrast, top-*k* sparsification substantially reduces density while preserving structured weight distributions.

**Table 1.** Mapping sparsity and entropy (sample 1-Pre).

| Assignment | $W_{\text{nnz}}$ | Sparsity (%) | $n_{\text{cells} \geq \epsilon}$ | Entropy |
| --- | --- | --- | --- | --- |
| Soft | 47,578,752 | 100 | 9,531 | 9.16 |
| Hard | 9,531 | 0.02 | 1 | 0.22 |
| Top-1% | 476,550 | 1.0 | 95 | 3.27 |
| Top-5% | 2,382,750 | 5.0 | 477 | 5.60 |
| Top-10% | 4,765,500 | 10.0 | 955 | 6.59 |

Downstream effects are shown in Table 2. TF ranking stability was evaluated using Spearman correlation relative to the soft baseline. Hard assignment substantially perturbs motif rankings (*ρ* = − 0.74), indicating near-inversion of regulatory ordering. Top-*k* sparsification reduces this deviation (*ρ* = −0.48 at 1%, *ρ* ≈ − 0.39 at 5–10%), demonstrating improved stability compared to hard assignment while retaining most projected ATAC signal.

**Table 2.** Projected ATAC retention and TF ranking stability (sample 1-Pre). TF rank correlation is computed relative to the soft assignment baseline.

| Assignment | Signal Retained (%) | TF Rank Corr. |
| --- | --- | --- |
| Soft | 100 | 1.0000 |
| Hard | 2.4 | -0.7365 |
| Top-1% | 41 | -0.4836 |
| Top-5% | 82 | -0.3953 |
| Top-10% | 95 | -0.3907 |

Top-*k* sparsification (5%) achieves an effective balance between computational efficiency (95% reduction in mapping density) and regulatory stability, and is therefore used as the default configuration in subsequent experiments.

### 3.3 Spatial Smoothness Analysis

Spatial regularization had limited impact on domain continuity and downstream TF ranking stability within the evaluated range (Table 3). Continuity remained high across settings, and TF rank concordance relative to the baseline was comparable, indicating that the mapping is relatively insensitive to moderate smoothness variation.

**Table 3.** Spatial smoothness ablation (sample 1-Pre). Continuity denotes the fraction of 6-NN spatial neighbor pairs sharing the same domain. TF Rank Corr. denotes the mean Spearman correlation of domain-enriched TF motif rankings across random seeds.

| $\lambda_{\text{cluster}}$ | Continuity | TF Rank Corr. |
| --- | --- | --- |
| 0 | $0.8484 \pm 0.0129$ | $-0.6583 \pm 0.0654$ |
| 0.5 | $0.8162 \pm 0.0059$ | $-0.7170 \pm 0.0560$ |
| 1 | $0.8165 \pm 0.0061$ | $-0.7297 \pm 0.0234$ |

### 3.4 Automated Spatial Domain Selection

Spatial domains were obtained via Leiden clustering on GraphST embeddings. Resolution was selected using a plateau-based heuristic balancing domain stability and silhouette score. The selected number of domains per section is reported in Table 4.

**Table 4.** Cohort summary (top-*k* 5% assignment). We define *n*_cells≥ε_ as the number of cells assigned to each spot with weight exceeding *ε* = 10^−4^.

| Sample | $n_{\text{spots}}$ | $n_{\text{cells}}$ | $n_{\text{genes}}$ | $n_{\text{domains}}$ | $W_{\text{nnz}}$ | $A_{\text{nnz}}$ | Mean $n_{\text{cells} > \varepsilon}$ |
| --- | --- | --- | --- | --- | --- | --- | --- |
| 1-Pre | 4,992 | 9,531 | 716 | 11 | 476,550 | 360,314,047 | 96 |
| 1-Post | 4,992 | 16,936 | 614 | 8 | 846,800 | 177,521,324 | 170 |
| 2-Pre | 4,992 | 16,391 | 470 | 7 | 819,550 | 236,766,896 | 164 |
| 2-Post | 4,992 | 15,577 | 870 | 8 | 778,850 | 185,202,745 | 156 |
| 3-Pre | 4,992 | 8,005 | 455 | 3 | 400,250 | 143,227,615 | 80 |
| 3-Post | 4,992 | 10,394 | 367 | 4 | 519,700 | 274,967,203 | 104 |
| 4-Pre | 4,992 | 9,791 | 680 | 8 | 489,550 | 149,888,149 | 98 |

### 3.5 Regulatory Inference and Concordance

Figure 1 illustrates spatial domains derived from (A) raw Visium clustering, (B) GraphST embedding, and (C) scRNA-projected spot domains on sample 1-Pre. The projected domains recapitulate large-scale tissue topology while preserving compartment boundaries, indicating that cross-modal mapping maintains spatial organization.

**Fig. 1.**
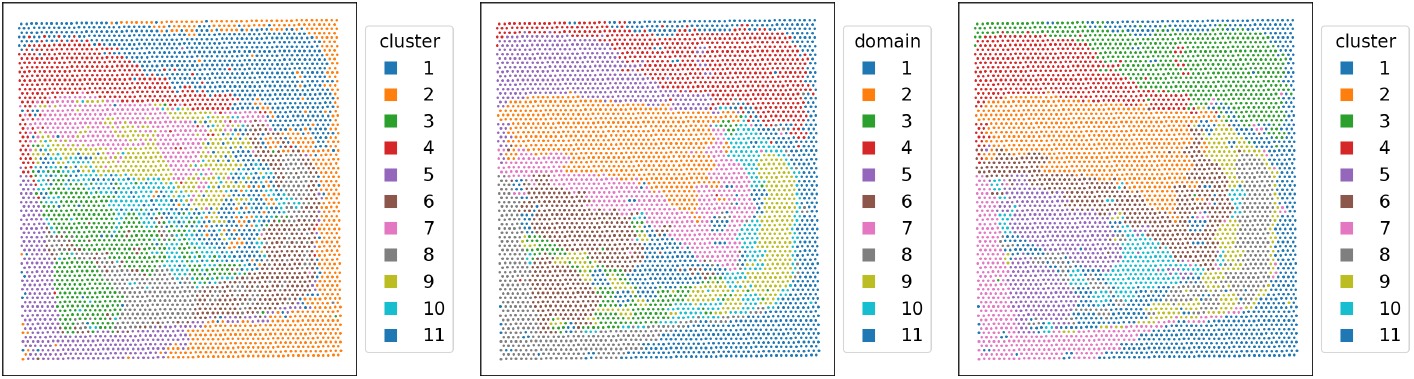
Spatial domain comparison on sample 1-Pre. (A) Source Visium spatial domains. (B) Domains inferred from GraphST embedding. (C) scRNA-projected spot domains. Spatial topology is preserved under projection, with coherent domain structure maintained across representations.

For each domain, we performed one-versus-rest differential expression (DE) and differential gene-proximal accessibility (DA; ±50 kb window including promoter regions). Regulatory concordance was quantified using Spearman correlation between domain-level DE and DA log fold changes across shared genes.

#### Example transcription factor enrichment

To illustrate spatial regulatory heterogeneity, we examined NFKB1 motif activity across domains (Table 5). NFKB1 exhibited strong enrichment in Domains 2 and 11 (logFC *>* 1.0, FDR *<* 10^−40^) and additional enrichment in Domains 3, 5, and 9, while Domains 1 and 4 showed significant depletion. These spatially restricted patterns are consistent with localized inflammatory regulatory programs rather than uniform activation across the tissue.

**Table 5.** Domain-specific NFKB1 motif enrichment. Positive logFC indicates enrichment relative to all other domains; negative logFC indicates depletion.

| Domain | logFC | FDR | Interpretation |
| --- | --- | --- | --- |
| 2 | 1.98 | $< 10^{-300}$ | Enriched |
| 3 | 1.09 | $1.0 \times 10^{-46}$ | Enriched |
| 5 | 0.28 | $8.2 \times 10^{-26}$ | Enriched |
| 9 | 0.57 | $3.0 \times 10^{-37}$ | Enriched |
| 11 | 1.14 | $2.9 \times 10^{-63}$ | Enriched |
| 1 | -0.98 | $2.9 \times 10^{-127}$ | Depleted |
| 4 | -1.33 | $4.4 \times 10^{-201}$ | Depleted |
| 6 | -0.41 | $8.2 \times 10^{-10}$ | Depleted |
| 7 | -0.57 | $1.2 \times 10^{-7}$ | Depleted |
| 8 | -0.88 | $6.1 \times 10^{-52}$ | Depleted |
| 10 | -0.43 | 0.57 | Not significant |

### 3.6 Cohort-Level Summary

Applying the default configuration (Top-5% assignment with automated domain selection) across seven sections demonstrates consistent sparsity, stable projected ATAC density, and reproducible domain discovery (Table 4).

## 4 Conclusion

We present a spatially informed multimodal integration framework for projecting single-cell transcriptomic and epigenomic profiles onto spatial transcriptomics data. Our analysis demonstrates that assignment strategy plays a central role in downstream regulatory inference. While dense soft mappings preserve signal, they are computationally expensive and difficult to interpret. In contrast, hard assignment severely degrades projected accessibility and destabilizes transcription factor ranking.

We show that top-*k* sparsification (5%) achieves an effective balance between sparsity and biological fidelity, retaining the majority of projected ATAC signal while preserving domain-level TF enrichment structure. Importantly, regulatory concordance between differential expression and gene-proximal accessibility remains consistent under the proposed mapping, indicating that biologically meaningful spatial regulatory patterns are preserved.

Across multiple melanoma sections, the framework produces stable spatial domains and reproducible regulatory signatures, including domain-specific enrichment of inflammatory regulators such as NFKB1. These results highlight the importance of assignment design in spatial multi-omics integration and provide a computationally efficient strategy for robust domain-level regulatory analysis.

Future work will extend the framework to larger cohorts, develop adaptive sparsification strategies to enhance biological stability under resource constraints, and perform comparative analyses of matched pre- and post-treatment samples.

## Acknowledgments

This project was supported by the National Center for Advancing Translational Sciences (NCATS), National Institutes of Health, through Grant Award Number UM1TR004539. The content is solely the responsibility of the authors and does not necessarily represent the official views of the NIH.

